# A Thermostable, Flexible RNA Vaccine Delivery Platform for Pandemic Response

**DOI:** 10.1101/2021.02.01.429283

**Authors:** Alana Gerhardt, Emily Voigt, Michelle Archer, Sierra Reed, Elise Larson, Neal Van Hoeven, Ryan Kramer, Christopher Fox, Corey Casper

## Abstract

Current RNA vaccines against SARS-CoV-2 are limited by instability of both the RNA and the lipid nanoparticle delivery system, requiring storage at −20°C or −70°C and compromising universally accessible vaccine distribution. This study demonstrates the thermostability and adaptability of a nanostructured lipid carrier (NLC) RNA vaccine delivery system for use in pandemic preparedness and pandemic response. Liquid NLC is stable at refrigerated temperatures for ≥ 1 year, enabling stockpiling and rapid deployment by point-of-care mixing with any vaccine RNA. Alternatively, NLC complexed with RNA may be readily lyophilized and stored at room temperature for ≥ 8 months or refrigerated temperature for ≥ 21 months. This thermostable RNA vaccine platform could significantly improve distribution of current and future pandemic response vaccines, particularly in low-resource settings.

**One Sentence Summary:** An RNA vaccine delivery system stable at room temperature for 8+ months and refrigerated for 21+ months.

## Main Text

RNA-based vaccines show great promise to effectively address existing and emerging infectious diseases (*1–3*), including the ongoing pandemic caused by the SARS-CoV-2 virus. RNA vaccines can be rapidly adapted to new targets and manufactured using sequence-independent operations, thus reducing the cost and time to develop new vaccines, particularly in pandemic settings (*4*). The recent Emergency Use Authorization granted to two safe and highly effective mRNA vaccines targeting SARS-CoV-2, less than one year after sequencing the novel coronavirus, highlights the power of this new technology (*5, 6*). However, one of the biggest challenges facing these extraordinary new vaccines is the ability to successfully distribute them widely in the face of a pandemic. Deep cold chain storage is required for both authorized vaccines (−70°C and −20°C for the SARS-CoV-2 RNA vaccines produced by Pfizer/BioNtech and Moderna, respectively). Frozen shipping and storage even at standard freezer conditions poses difficulties in settings with well-established medical infrastructure – challenges greatly compounded in areas with limited resources (*7–9*).

Lack of stability in RNA vaccines is a critical issue, but the physicochemical reasons behind this are under-studied and poorly understood (*9*). However, several facts are clear. First, vaccine RNA molecules are prone to cleavage by ubiquitous ribonucleases (i.e. RNases). Engineering of the RNA has previously been done in order to stabilize it, as reviewed in (*10*), but stability problems remain. Second, due to its size, negative charge, and hydrophilicity, RNA alone cannot easily cross a cell membrane to enter target cells upon injection (*11*). Thus, RNA delivery formulations are needed to stabilize and protect RNA molecules from degradation (reviewed in (*12, 13*)). The current system of choice for delivering RNA vaccines, including all SARS-CoV-2 vaccines in clinical trials to date, is a lipid nanoparticle (LNP) delivery system (*5, 14–17*) in which the negatively-charged RNA molecule is encapsulated within a multicomponent lipid system. This results in 70-100 nm diameter RNA/LNP complexes which protect the RNA from RNase degradation and allow for successful endocytosis by the cell (reviewed by (*18, 19*)). However, stability of both the RNA and LNP remain an issue (*9*), with sensitivity to frozen temperatures resulting in detrimental impacts to their colloidal stability after freeze/thaw (*20, 21*).

A number of alternative lipid-based delivery systems have been proposed and developed to deliver RNA vaccines (*22–24*); however, a critical current need remains an effective and thermostable RNA vaccine delivery system (*9*). We demonstrate the ability of a lyophilizable, thermostable nanostructured lipid carrier (NLC) to effectively deliver both mRNA- and replicating RNA-based vaccines by intramuscular injection. The liquid NLC alone maintains stability for at least 1 year of storage at refrigerated temperatures while lyophilized NLC/RNA complexes are shown to retain biophysical properties and ability to induce protein expression in vivo after at least 8 months of room temperature storage and at least 21 months at refrigerated temperatures.

The NLC delivery system consists of an oil core comprised of solid (trimyristin) and liquid (squalene) lipids surrounded by surfactants (sorbitan monostearate and polysorbate 80) and a cationic lipid (DOTAP) (*25*). RNA complexes electrostatically to the outside of an NLC particle (Figure 1A). The NLC system itself displays long-term stability at 4°C, maintaining its particle size and component concentrations (Figure 1B, 1C), as well as retaining its ability to complex with and protect RNA from RNase challenge (Figure 1D, 1E). Due to this long-term stability, NLC is suitable for stockpiling for pandemic preparedness applications – RNA targeting a specific pathogen can be rapidly produced in response to a pandemic and complexed with pre-manufactured and stockpiled NLC.

**Fig 1.**
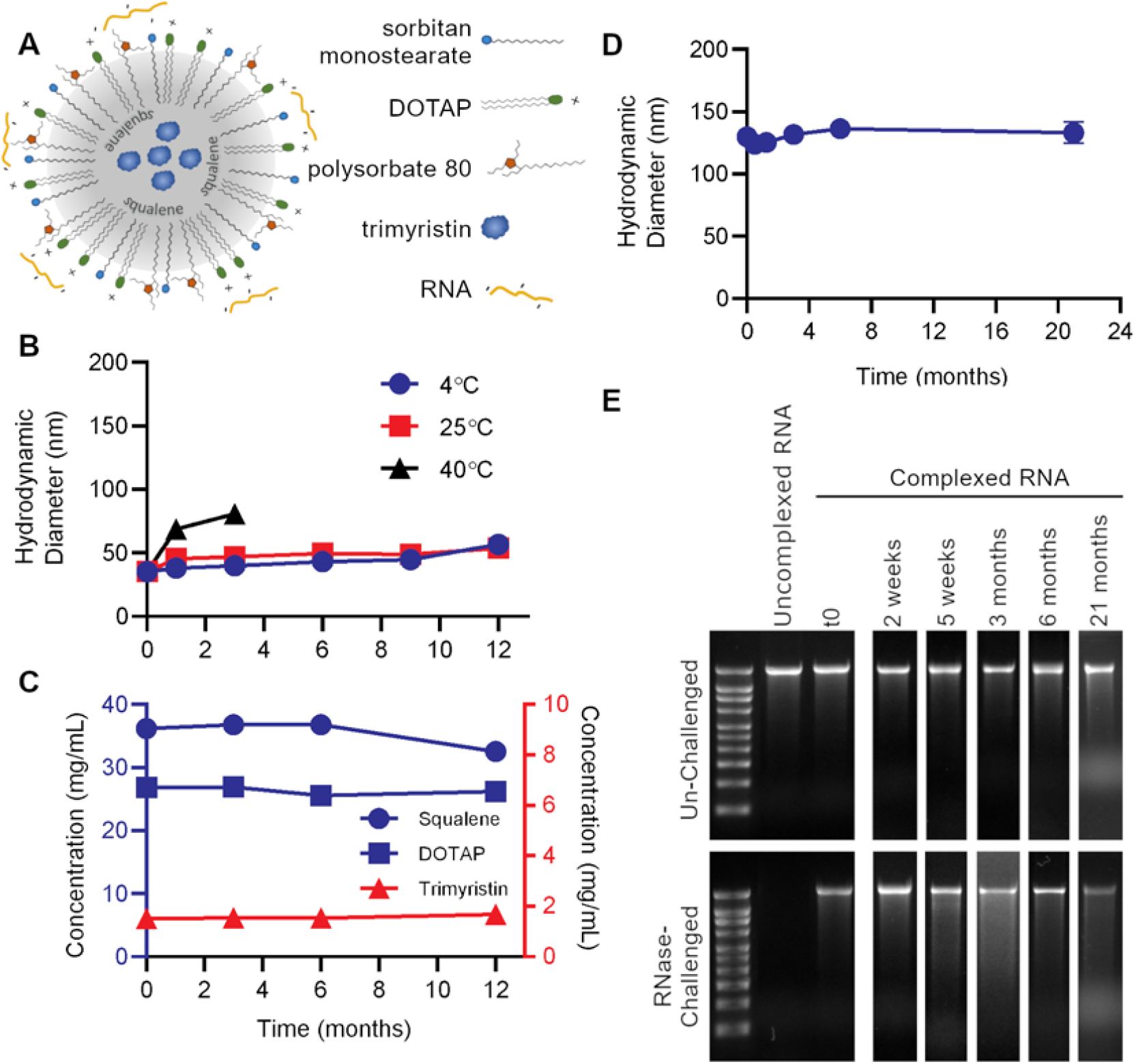
Nanostructured lipid carrier (NLC) formulation alone is stable at 4°C, allowing for stockpiling. (A) Schematic of RNA electrostatically binding to the outside of the NLC. (B) Particle size of NLC alone after storage at indicated temperatures. (C) Concentration of NLC components after long-term 4°C storage. (D) Vaccine particle size after complexing 4°C-stored NLC with SEAP saRNA. (E) Protection of SEAP saRNA from RNase challenge by NLC stored at 4°C for the indicated length of time. Full gel images are in Supplemental Figure S4.

Beyond its utility for stockpiling, NLC complexed with RNA is a thermostable vaccine platform which can greatly ease the challenges of distributing RNA vaccines in both pandemic and non-pandemic situations. We previously demonstrated the utility of an NLC/self-amplifying RNA (saRNA) vaccine against Zika virus that induced high levels of neutralizing antibodies and protected mice against viral challenge (*23*). Here, we demonstrate that the next generation of this Zika NLC/saRNA vaccine (Supplemental Figure S1A) can be successfully lyophilized for long-term storage (Figure 2) with the addition of 10% w/v sucrose as a lyoprotectant. The presence of sucrose promotes the formation of a dense, white, lyophilized cake (Supplemental Figure S2) and also serves to protect the components of the system against the stresses encountered during freezing, drying, and reconstitution.

**Fig. 2.**
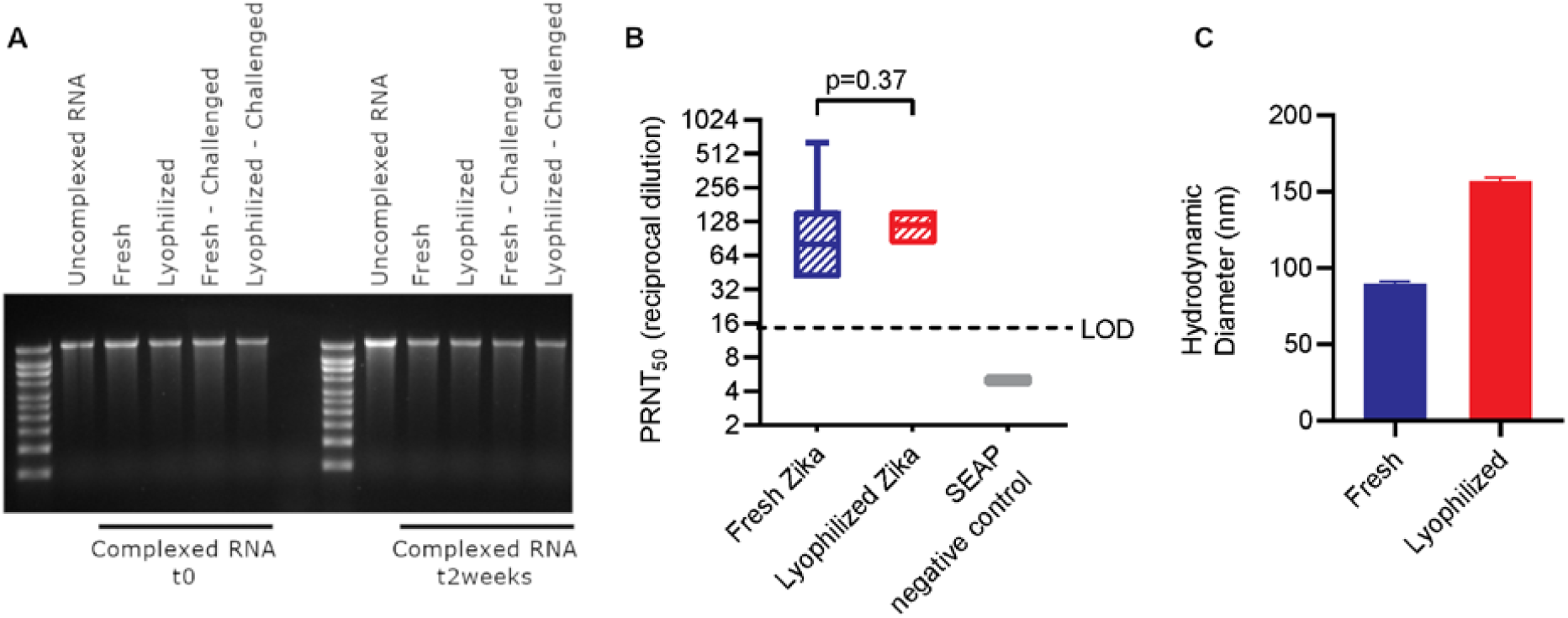
Comparison of lyophilized Zika NLC/saRNA vaccine with freshly complexed vaccine. (A) Integrity of Zika saRNA under fresh or lyophilized/reconstituted conditions after it has been extracted from the NLC and protection of Zika saRNA after it has been challenged with RNase and then extracted from the NLC (“Challenged”). The fresh and lyophilized/reconstituted vaccine were also evaluated under un-challenged and challenged conditions after 2 weeks of storage at 4°C. (B) in vivo immunogenicity equivalence of fresh and lyophilized/reconstituted Zika vaccine by PRNT. SEAP NLC/saRNA was used as an in vivo negative control. n=10 mice in all groups. (C) Hydrodynamic diameter of fresh and lyophilized/reconstituted vaccine by Dynamic Light Scattering (DLS).

RNA integrity and protection against RNase challenge is maintained after lyophilization/reconstitution as shown by agarose gel electrophoresis (*25*) of RNA extracted from NLC/RNA (Figure 2A). Furthermore, both the freshly-complexed liquid and the lyophilized/reconstituted vaccines are stable for at least two weeks at refrigerated temperatures (Figure 2A), retaining their ability to protect from RNase challenge as compared to both freshly mixed and freshly reconstituted lyophilized vaccine. Upon reconstitution and i.m. injection into C57BL/6 mice, the lyophilized Zika saRNA vaccine is able to induce neutralizing (Figure 2B) antibody titers identical to freshly-complexed, un-lyophilized vaccine at the same 1 μg dose. The size of the complex has a moderate increase post-lyophilization and reconstitution (Figure 2C), which does not appear to affect in vivo efficacy.

As mRNA-based vaccines have been the frontrunners in Covid-19 response, the ability of a thermostable delivery system to effectively deliver vaccine mRNA is a critical need now and in the future. The flexibility and utility of this NLC-based system is shown by complexing it with commercially-available mRNA encoding ovalbumin (OVA). Biophysical characterization (*25*) of the NLC/mRNA complexes shows nearly identical protection of the mRNA against Rnase challenge (Figure 3A) as with Zika NLC/saRNA complexes. Furthermore, by increasing the concentration of the lyoprotectant sucrose to 20% w/v, the particle size exhibits only a slight increase post-lyophilization and reconstitution (Figure 3B), similar to that observed in a complex that is frozen. Images of lyophilized OVA NLC/mRNA are in Supplemental Figure S3.

**Fig. 3.**
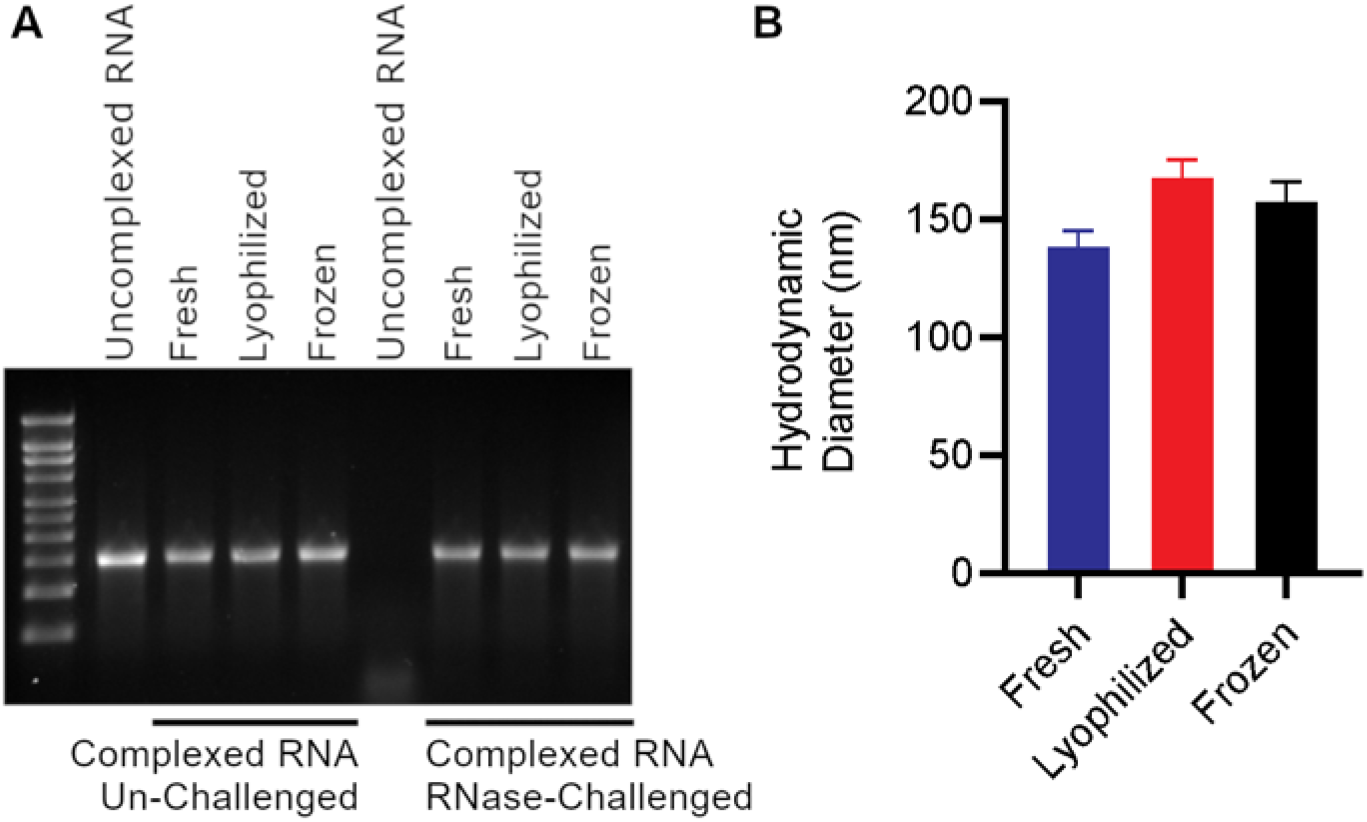
Comparison of lyophilized or frozen OVA NLC/mRNA with freshly complexed material. (A) Integrity of OVA mRNA under fresh, frozen, or lyophilized conditions after it has been extracted from the NLC complex (“Un-Challenged”) and protection of OVA mRNA after it has been challenged with RNase and then extracted from the NLC complex (“Challenged”). (B) Hydrodynamic diameter of fresh, frozen, and lyophilized complexes by Dynamic Light Scattering (DLS).

Finally, we demonstrate the long-term thermostability of the NLC-based RNA vaccine platform using a self-amplifying RNA antigen expression reporter system expressing secreted alkaline phosphatase (SEAP-saRNA), which allows for sensitive mouse serum detection of i.m.-injected saRNA (*25*). Lyophilized SEAP NLC/saRNA complexes with 20% w/v sucrose as a lyoprotectant stored at 4°C, 25°C, and 40°C are compared with frozen complexes stored at −80°C and −20C°, liquid complexes stored at 4°C and 25°C, and freshly made complexes prepared each analysis day. All lyophilized samples maintain an elegant, white cake throughout the study with no discoloration or cracking and minimal cake shrinkage. All lyophilized samples readily reconstitute with nuclease-free water, and the reconstituted complexes are visually similar to freshly-complexed comparators (Figure 4A).

**Fig. 4.**
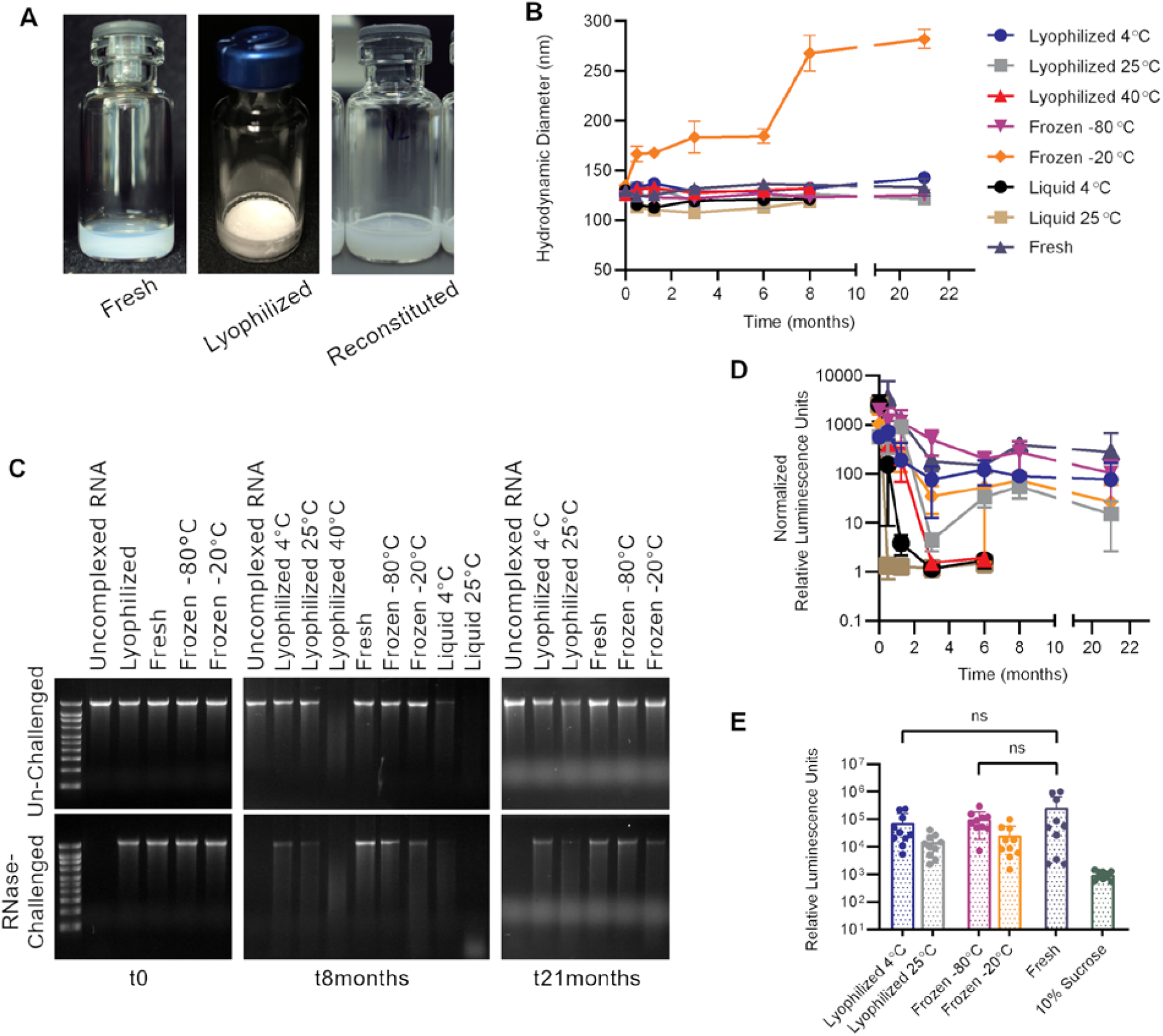
SEAP NLC/saRNA under lyophilized, frozen, or liquid storage conditions in comparison to freshly complexed material. (A) Vial images of freshly complexed, lyophilized, and reconstituted material at t0. (B) Hydrodynamic diameter of the complexes over time as compared to a freshly complexed control. (C) RNA integrity of the stored samples by agarose gel electrophoresis at t0, t8months, and t21months and protection from RNase challenge at each timepoint. Gel images at all timepoints are shown in Supplemental Figure S3. (D) Normalized in vivo SEAP expression for lyophilized, frozen, or liquid stored samples in comparison with freshly complexed material after long-term storage. (E) Comparable in vivo SEAP expression at 21 months for lyophilized vaccine stored at 4°C, frozen vaccine stored at −80°C, and freshly-prepared vaccine; 10% sucrose group shown as control.

Initially, all NLC/saRNA complexes (Figure 4B) measure 125±10 nm in diameter, including liquid, frozen, and lyophilized versions. Differences of less than 15% are observed between the initial and final timepoints for all conditions except frozen material stored at −20°C. This demonstrates the excellent colloidal stability of NLC/RNA complexes, allowing them to withstand the stresses of the lyophilization process and long-term storage, even at elevated temperatures (40°C for lyophilized and 25°C for liquid storage). It is interesting to note that, while size stability is not maintained for complexes stored at −20°C, this did not impact the ability of the NLC/saRNA complex to express protein in vivo.

RNA integrity and protection from RNase challenge is demonstrated by agarose gel electrophoresis after extraction from the stored NLC complexes (Figure 4C), and the ability of stored NLC/saRNA to express protein in vivo is demonstrated by injection of 100 ng i.m. into C57BL/6 mice with assay of SEAP expression in sera 5 days post-injection relative to sucrose-injected mouse sera (Figure 4D). RNA integrity in the NLC/saRNA complexes is again maintained after lyophilization and after freeze/thaw. After 8 months of storage, lyophilized complex stored at 4°C and 25°C and complex stored frozen at −80°C and −20°C show comparable levels of mouse serum SEAP expression to the freshly complexed material. Furthermore, lyophilized material maintains RNA integrity, protection against RNase challenge, and in vivo expression ability for at least 21 months when stored at refrigerated (4°C) temperatures. Under accelerated conditions, degradation in the form of reduced protection from RNase challenge and loss of in vivo protein expression is observed at 2 weeks for the liquid 25°C condition, at 5 weeks for the liquid 4°C condition, and at 3 months for the lyophilized 40°C condition (Supplemental Figure S4).

RNA vaccines are important tools to combat existing and emerging infectious diseases, including SARS-CoV-2, due to their rapid adaptability to new target pathogens (*1–6*). However, strict cold chain requirements for current RNA vaccine formulations greatly complicate global distribution and increase cost, leading to calls for rapid advances in the stability of RNA vaccine formulations (*9*). We demonstrate that a safe and effective NLC-based RNA vaccine delivery system (*23*) has greatly increased thermostability over current LNP formulations. The liquid NLC alone is stable at refrigerated temperatures for at least two years, and NLC complexed with mRNA or saRNA is able to be stored in lyophilized, liquid, and frozen forms for extended periods of time. Moreover, upon reconstitution, NLC-formulated RNA vaccine retains its integrity for at least 2 weeks at refrigerated temperatures. This NLC-based delivery technology represents a significant advance for RNA vaccines with potentially paradigm-shifting implications on vaccine manufacture, storage, distribution, and overall cost due to its thermostable properties.

We hypothesize multiple mechanisms behind the improved thermostability of NLC-based delivery formulations relative to LNP-based formulations. First, the robust physical stability of the NLC allows for minimal growth in particle size, retention of constituent components, and maintenance of complexing compatibility for at least one year under refrigerated storage. Furthermore, the NLC system provides excellent protection to the RNA against RNases, presumably due to the electrostatic interaction between RNA’s negatively-charged phosphate backbone and the positively-charged amine group of the NLC’s DOTAP component. This interaction drives RNA/NLC complex formation and protects the RNA from cleavage by RNases during long-term storage and after administration.

Most importantly, the physical characteristics of this NLC-based RNA vaccine formulation allow for lyophilization, a technique commonly used to stabilize vaccines and biologics and eliminate a deep cold chain requirement (*7, 8, 26–29*). In lyophilized drug products, non-reducing sugars (such as sucrose) act as lyoprotectants through multiple proposed mechanisms such as replacing water in hydrogen bonding with the components of the system or enclosing the system within the rigid sugar matrix of the dried state where enzymatic or other degradation is limited (*30*). While lyophilization of liposome-based formulations has been attempted for decades (reviewed in (*30*)), it is notoriously difficult due to the liposome’s physical structure (i.e. a lipid bilayer surrounding a core aqueous phase) which is disrupted by the freezing and drying steps of lyophilization. Recent published attempts at LNP/RNA complex lyophilization have been semi-successful at best, showing significant loss of RNA activity despite the addition of lyoprotectants (*20, 21*). While optimization of LNP lyophilization may yet be attempted (reviewed in (*31*)), the technical challenge of redesigning and clinically testing lyophilizable liposome-based RNA vaccine delivery formulations is significant and without guaranteed success.

The NLC system is ideal for situations of pandemic response. NLC manufacture is straightforward and scalable since it employs similar processes and equipment as oil-in-water emulsion technology already employed in licensed vaccines – properties essential to best support large-scale pandemic response. For pandemic preparedness, the long-term refrigerator-stable NLC alone could be stockpiled to enable rapid response. Furthermore, as RNA of different lengths or with multiple genetic variations can be rapidly synthesized and complexed on the outside of the NLC, head-to-head comparisons of different RNA species is feasible and such a vaccine may be rapidly adapted to evolving viral variants or emerging pathogens. Finally, once an RNA vaccine candidate has been chosen, the potential for a lyophilized, heat-stable RNA vaccine drug product would maximize the speed of global vaccine distribution.

We do note that the presented long-term stability data of the lyophilized system is with RNA expressing a reporter protein rather than any vaccine antigen. However, it is both highly likely that SARS-CoV-2 vaccine antigen expression would be similar to that of the reporter protein expression and obvious that insufficient time has elapsed since the start of the pandemic for such long-term stability data to be available on any Covid-19 antigen-specific vaccine. Additionally, long-term stability of the NLC formulation alone is relevant for application to any vaccine target. Finally, while this specific NLC-based formulation has not yet been clinically tested, safety concerns are low. Squalene, polysorbate 80, and sorbitan monostearate are already in FDA-licensed drug products, and trimyristin is commonly used in the cosmetics industry and closely related to tristearin, found in licensed drug products. Finally, DOTAP has been successfully evaluated in multiple clinical trials (*32*). We have injected NLC/saRNA complexes at doses up to 800 μg saRNA into NHPs with no signals of reactogenicity or toxicity.

While the current study effectively demonstrates the excellent thermostability of an NLC/RNA drug product, further work should be conducted to push the limits of storage and use conditions. The excellent thermostability achieved here was a proof-of-concept attempt for this system. It is likely that further improvement to the lyophilized RNA/NLC drug product is possible, through optimization of excipient concentrations and lyophilization cycle parameters. Optimization may allow for additional enhancement of RNA vaccine drug product thermostability, allowing retention of bioactivity under extreme temperatures and/or temperature cycling conditions that may be experienced for a globally-distributed vaccine.

## Acknowledgments

The authors would like to thank Jacob Archer, Julie Bakken, Peter Battisti, Stacey Ertel, Jasmine Fuerte-Stone, Brian Granger, and Jasneet Singh for technical assistance.

## Author contributions

Conceptualization – A.G., E.V., M.A., N.V.H., R.K., C.C.; Methodology - A.G., E.V., S.R., E.L., M.A.; Investigation – A.G., E.V., S.R., E.L., M.A.; Writing – Original Draft – A.G., E.V.; Writing – Review & Editing – A.G., E.V., S.R., E.L., M.A., R.K., C.F., C.C.; Supervision – A.G., E.V., N.V.H., R.K., C.F., C.C.; Funding acquisition – N.V.H., C.C.

## Data and materials availability

All data is available in the main text or the supplementary materials.

## Supplementary Materials for

### Materials and Methods

#### saRNA DNA Templates

DNA templates for self-amplifying RNA (saRNA) encoding the Zika pre-membrane (PrM) and envelope (E) proteins were produced as previously described (*23*). Briefly, sequences for the Zika virus signal peptide at the N-terminal end of the capsid protein through the prM and E genes were taken from ZIKV strain H/PF/2013 (GenBank Accession #KJ776791), codon-optimized for mammalian expression, and subcloned into a T7-TC83 plasmid. The resulting plasmid pT7-VEE-Zika-prME contains the 5’ UTR, 3’ UTR, and non-structural proteins derived from the attenuated TC-83 strain of VEEV, with the aforementioned Zika virus genes replacing the VEEV structural proteins downstream of a subgenomic promoter (Supplemental Figure S1A). Plasmid pT7-VEE-Zika-prME varies slightly from the previously-published Zika vaccine plasmid (*23*) with a change of the antibiotic resistance gene from Ampicillin to Kanamycin to allow for GMP manufacture and an optimization of the subgenomic promoter for antigen expression enhancement.

Similarly, DNA templates for self-amplifying RNA encoding the secreted alkaline phosphatase protein (SEAP) were constructed in two different versions (Supplemental Figure S1B, S1C). The first, pT7-VEE-SEAP-V1, is identical to that published in (*23*) and was used as the template for all SEAP-saRNA used in the long-term stability studies shown in Figure 4. An updated version (pT7-VEE-SEAP-V2) reflects the same antibiotic resistance gene and subgenomic promoter changes described above to allow for optimal comparison to pT7-VEE-Zika-prME in the vaccine immunogenicity studies in Figure 2. All plasmid sequences were confirmed using Sanger sequencing. DNA templates were amplified in *E. coli* and isolated using maxi or gigaprep kits (Qiagen) and linearized by NotI restriction digest (New England Biolabs). Linearized DNA was purified by phenol chloroform extraction.

#### RNA Production and Purification

Generation of saRNA stocks was achieved by T7 promoter-mediated in vitro transcription using NotI-linearized DNA template. In vitro transcription was performed using an in house-optimized protocol using T7 polymerase, RNase inhibitor, and pyrophosphatase enzymes procured from Aldevron. DNA plasmid was digested away (DNase I, Aldevron) and cap0 structures were added to the transcripts by vaccinia capping enzyme, GTP, and S-adenosyl-methionine (Aldevron). RNA was then purified from the transcription and capping reaction components by chromatography using a CaptoCore 700 resin (GE Healthcare) followed by diafiltration and concentration using tangential flow filtration. The saRNA material was terminally filtered with a 0.22µm polyethersulfone filter and stored at −80°C until use. All saRNA was characterized by agarose gel electrophoresis and quantified both by UV absorbance (NanoDrop 1000) and Ribogreen assay (Thermo Fisher). Ovalbumin-expressing mRNA was obtained from a commercial vendor (TriLink CleanCap OVA mRNA, L-7610).

#### NLC Formulation Production

The nanostructured lipid carrier (NLC) formulation was prepared as described previously (*23*). Briefly, squalene (Sigma), sorbitan monostearate (Sigma), DOTAP (Corden), and trimyristin (IOI Oleochemical) were mixed and heated at 70°C in a bath sonicator. Separately, polysorbate 80 (Fisher Scientific) was diluted in 10 mM sodium citrate trihydrate and also heated to 70°C in a bath sonicator. After all components were dissolved, the oil and aqueous phases were mixed at 7,000 rpm in a high-speed laboratory emulsifier (Silverson Machines). The mixture was then processed by high-shear homogenization to further decrease particle size. Using an M-110P microfluidizer (Microfluidics), the colloid mixture was processed at 30,000 psi for eleven discrete microfluidization passes. The NLC product was terminally filtered with a 0.22µm polyethersulfone filter and stored at 2°C–8°C until use.

#### NLC Formulation Component Assay

The concentrations of DOTAP, squalene, and trimyristin in the NLC were determined by High Performance Liquid Chromatography (HPLC). Samples were prepared in triplicate, diluted 1:20 in HPLC mobile phase B (50 µL sample into 950 µL mobile phase B), injected at 10 µL injection volume, then analyzed using an Agilent 1100 quaternary pump HPLC system in combination with a Corona Veo charged aerosol detector (CAD). The method utilized a Phenomenex Synergi Hydro RP C18 80 A column (4 µm 4.6 × 250 mm) with a two solvent system gradient consisting of a mixture of 75:15:10 methanol:chloroform:water (mobile phase A) and a 1:1 mixture of methanol:chloroform (mobile phase B), with both mobile phases containing 20 mM ammonium acetate and 1% acetic acid. The system was held at 35°C and run at a flow rate of 1 mL/min. DOTAP, trimyristin, and squalene were dissolved in mobile phase B, and the injection volume was varied to create a 5-point standard curve.

#### NLC/RNA Complexing

NLC/RNA complexes were prepared at a nitrogen:phosphate (N:P) ratio of 15 for all cases. Fresh complexes were prepared by mixing RNA 1:1 by volume with NLC prepared in a buffer containing 10 mM sodium citrate and 20% w/v sucrose to achieve a final complex containing 200 ng/μL RNA in an isotonic 5 mM sodium citrate and 10% w/v sucrose aqueous buffer. Complexes for lyophilization were prepared with 10% or 20% w/v sucrose as noted in the text without additional sodium citrate. Complexes were incubated on ice for 30 minutes after mixing to ensure complete complexing.

#### NLC/RNA Complex Lyophilization

Lyophilized complex was prepared using a Virtis AdVantage 2.0 EL-85 bench-top freeze dryer controlled by the microprocessor-based Wizard 2.0 software. The lyophilization cycle consisted of a freezing step at −50°C, a primary drying step at −30°C and 50 mTorr, and a secondary drying step at 25°C and 50 mTorr. At the completion of the cycle, samples were brought to atmospheric pressure, blanketed with high purity nitrogen, and stoppered prior to being removed from the freeze-dryer chamber. Lyophilized material was reconstituted using nuclease-free water and gently swirled. Reconstituted material was diluted to 5 mM sodium citrate and 10% w/v sucrose (for isotonicity) prior to any in vivo experiments.

#### Particle Size Characterization

Hydrodynamic diameter (particle size) of both the NLC formulation alone and the NLC/RNA complex was determined using dynamic light scattering (Zetasizer Nano ZS, Malvern Instruments). Samples were diluted 1:100 in nuclease-free water in triplicate preparations and measured in a disposable polystyrene cuvette with the following parameters: material RI = 1.59, dispersant RI (water) = 1.33, T = 25^°^C, viscosity (water) = 0.887 centipoise [cP], measurement angle = 173° backscatter, measurement position = 4.65 mm, automatic attenuation.

#### NLC/RNA Complex RNase Protection Assay

Integrity of RNA after complexing and protection against RNase challenge was evaluated by agarose gel electrophoresis. Fresh, frozen/thawed, or lyophilized/reconstituted samples were diluted to a final RNA concentration of 40 ng/μL in nuclease-free water. For RNase-challenged samples, the diluted RNA was incubated with RNase A (Thermo Scientific) for 30 minutes at room temperature at amounts sufficient to completely degrade un-complexed RNA (ratios of 1:40 RNase:SEAP-RNA and 1:200 RNase:Zika-RNA). This was followed by treatment with recombinant Proteinase K (Thermo Scientific) at a ratio of 1:100 RNase A:Proteinase K for 10 minutes at 55°C. For both challenged and un-challenged samples, RNA was then extracted from the complexes by adding 25:24:1 phenol:chloroform:isoamyl alcohol (Invitrogen) to the complex 1:1 by volume, vortexing, and centrifuging at 17,000*g* for 15 minutes. The supernatant for each sample was mixed 1:1 by volume with Glyoxal load dye (Invitrogen) and incubated at 50°C for 20 minutes. For each complex, 200 ng of RNA was loaded and run on a denatured 1% agarose gel at 120 V for 45 minutes in Northern Max Gly running buffer (Invitrogen). Un-complexed RNA under challenged and un-challenged conditions was included in each gel as a control. Millenium RNA marker (ThermoFisher) was included on each gel with markers at 0.5, 1, 1.5, 2, 2.5, 3, 4, 5, 6, and 9 kilobases. Gels were imaged using ethidium bromide protocol on a ChemiDoc MP imaging system (BioRad).

#### Mouse Studies

C57BL/6J mice between 4 and 8 weeks of age at study onset obtained from The Jackson Laboratory were used for all animal studies in this work. All animal work was done under the oversight of IDRI’s Institutional Animal Care and Use Committee and/or the Bloodworks Northwest Research Institute’s Institutional Animal Care and Use Committee. All animal work was in compliance with all applicable sections of the Final Rules of the Animal Welfare Act regulations (9 CFR Parts 1, 2, and 3) and the NIH *Guide for the Care and Use of Laboratory Animals: Eighth Edition*.

#### Zika NLC/saRNA *in vivo* Immunogenicity

To compare immunogenicity of lyophilized/reconstituted versus freshly-complexed Zika NLC/saRNA vaccines, mice (n=10/group) were immunized with 1 μg of freshly-complexed Zika NLC/saRNA vaccine, 1 μg lyophilized/reconstituted Zika NLC/saRNA vaccine, or 10 μg of SEAP NLC/saRNA complex as a negative control. Vaccine was injected intramuscularly in 50 μl volumes in both rear quadriceps muscles of each mouse for a total of 100 ul vaccine per mouse. Injections sites were monitored for signs of reactogenicity for the 3 days post-injection, with no such signs noted. Blood samples were taken from all immunized mice 14 days post-immunization by the retro-orbital route for serum antibody assays by PRNT.

#### ZIKV PRNT

PRNT assays were performed on mouse serum samples to quantify neutralizing antibody titers. Briefly, Vero (ATCC CCL-81) cells were cultured at standard conditions (37°C, 5% CO2) in antibiotic-free high-glucose DMEM supplemented with GlutaMax (Gibco) and 10% v/v heat-inactivated FBS (HyClone). Cells were plated at a density of 5 × 10^5^ cells/well in 6 well plates (Corning) and incubated overnight to form 90% confluent monolayers. Mouse serum samples were serially diluted 1:2 in DMEM containing 1% heat-inactivated FBS. All serum dilutions were then diluted 1:2 with 100 PFU of ZIKV strain FSS13025 and incubated at 37°C for 1 hr. Cell supernates were removed and replaced with 200 ul of the virus/serum dilutions and allowed to incubate at culture conditions for 1 hour with gentle rocking every 20 minutes. Two ml of overlay medium comprised of DMEM containing 1% agarose (SeaKem), GlutaMax, and 1% v/v FBS was added to each well, allowed to solidify, and plates were incubated for 3 days at standard culture conditions. Cells were then fixed in 10% formalin (Fisher Scientific) for 20 minutes and stained with crystal violet for plaque visualization and counting.

#### In vivo Functionality of Stored SEAP NLC/saRNA

To verify the in vivo functionality of long-term stored SEAP NLC/saRNA complexes, mice (n=5 for t0 to t8 months and n=10 for t21 months) received a total dose of 100 ng RNA in a single 50 μL i.m. injection in one hind leg. A control group of mice received a 50 μL i.m. injection of 10% sucrose in a hind leg. Blood samples were taken from all immunized mice on days 3, 5, and 7 post-injection by the retro-orbital route. Serum samples were assayed for SEAP expression using the NovaBright Phospha-Light EXP Assay Kit for SEAP (ThermoFisher) according to the manufacturer’s directions. Relative luminescence was measured using a Biotek Synergy2 plate reader. At each timepoint, SEAP expression for sample at each storage condition was normalized to the SEAP expression of the 10% sucrose control.

#### Statistical Analyses

Comparability of PRNT titers between lyophilized and freshly complexed vaccine presentations for the saRNA Zika vaccine (Figure 2B) were conducted by a 2-tailed homoscedastic t-test on natural log-transformed PRNT titers. Log-transformed data were visually assessed for normality prior to analysis. Comparability of SEAP expression levels at t21months for each stored sample to a freshly complexed control (Figure 4E) was conducted using Dunnett’s multiple comparisons test on the data prior to normalization.

**Fig. S1.**
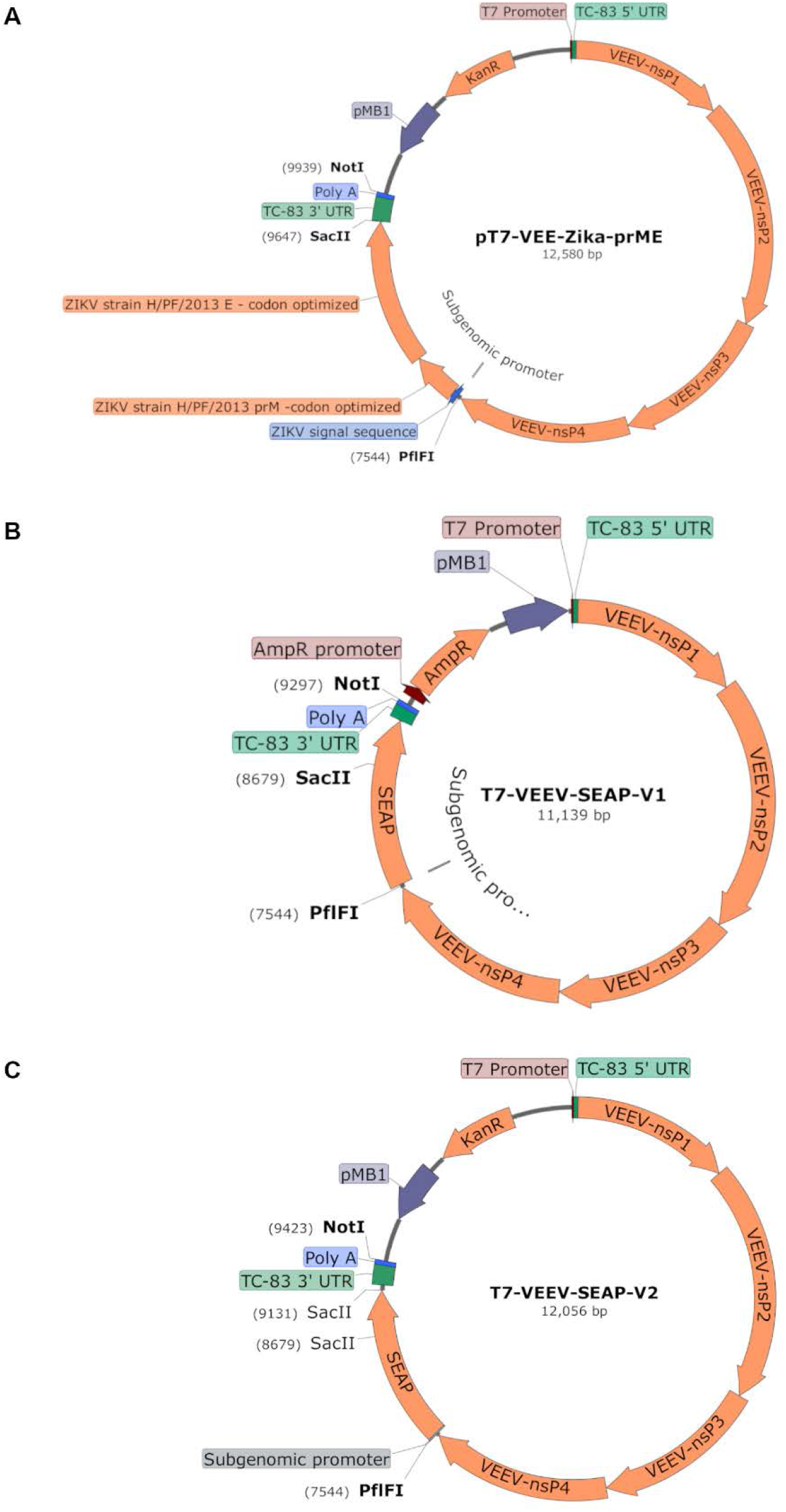
DNA plasmid templates for self-amplifying RNA vaccine constructs. RNA replicons consist of the 5’UTR, non-structural proteins, and 3’UTR sequences of the attenuated TC-83 strain of Venezuelan equine encephalitis virus (VEEV), with Zika virus PrM-E genes (A), or the secreted alkaline phosphatase (SEAP) gene (B,C) inserted in place of VEEV structural proteins, as described in the section “saRNA DNA Templates”.

**Fig. S2.**
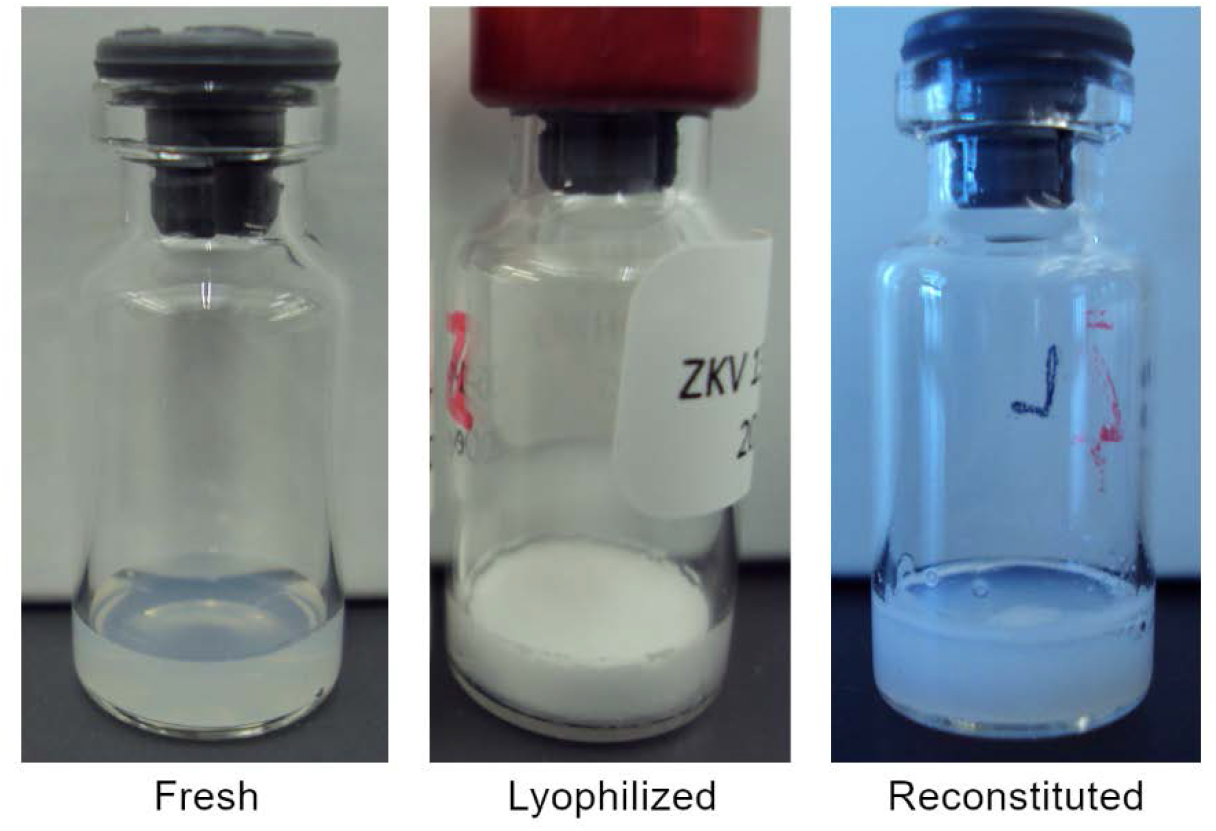
Vial images of fresh, lyophilized, and reconstituted Zika NLC/saRNA vaccine.

**Fig. S3.**
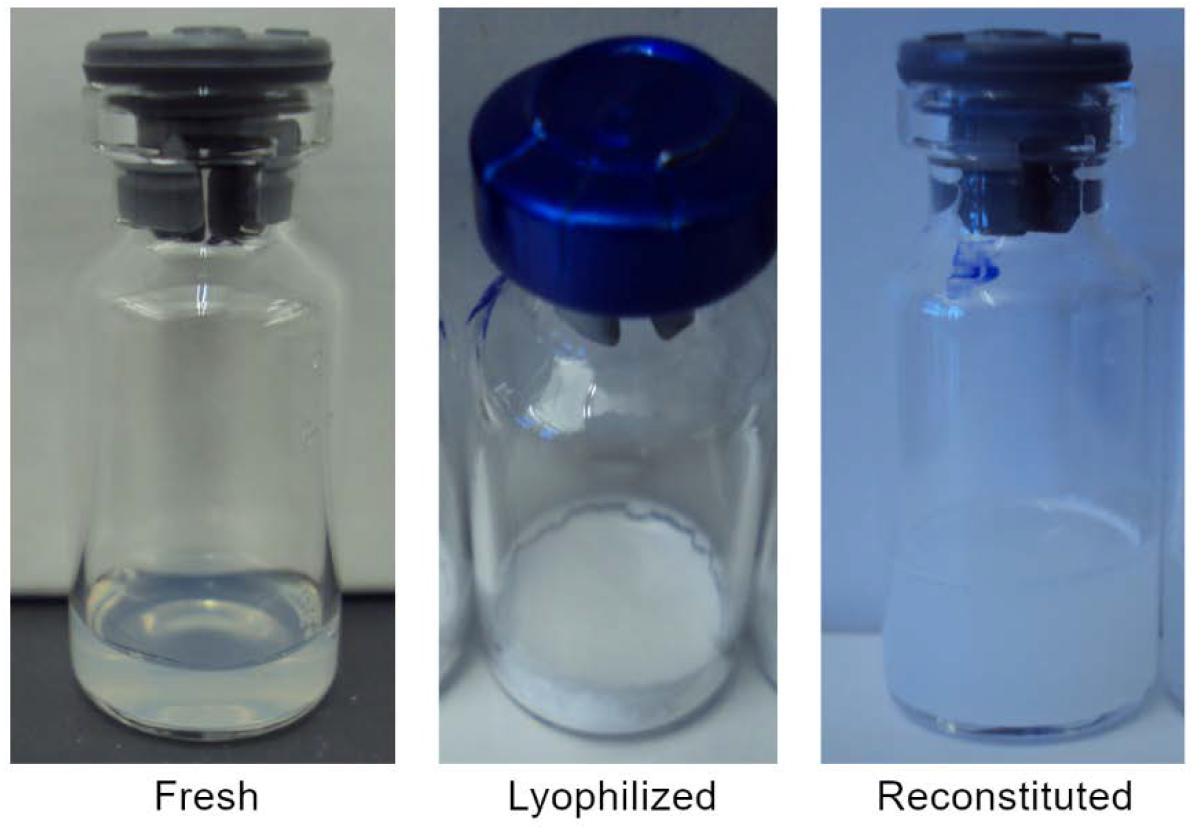
Vial images of fresh, lyophilized, and reconstituted OVA NLC/mRNA complex.

**Fig. S4.**
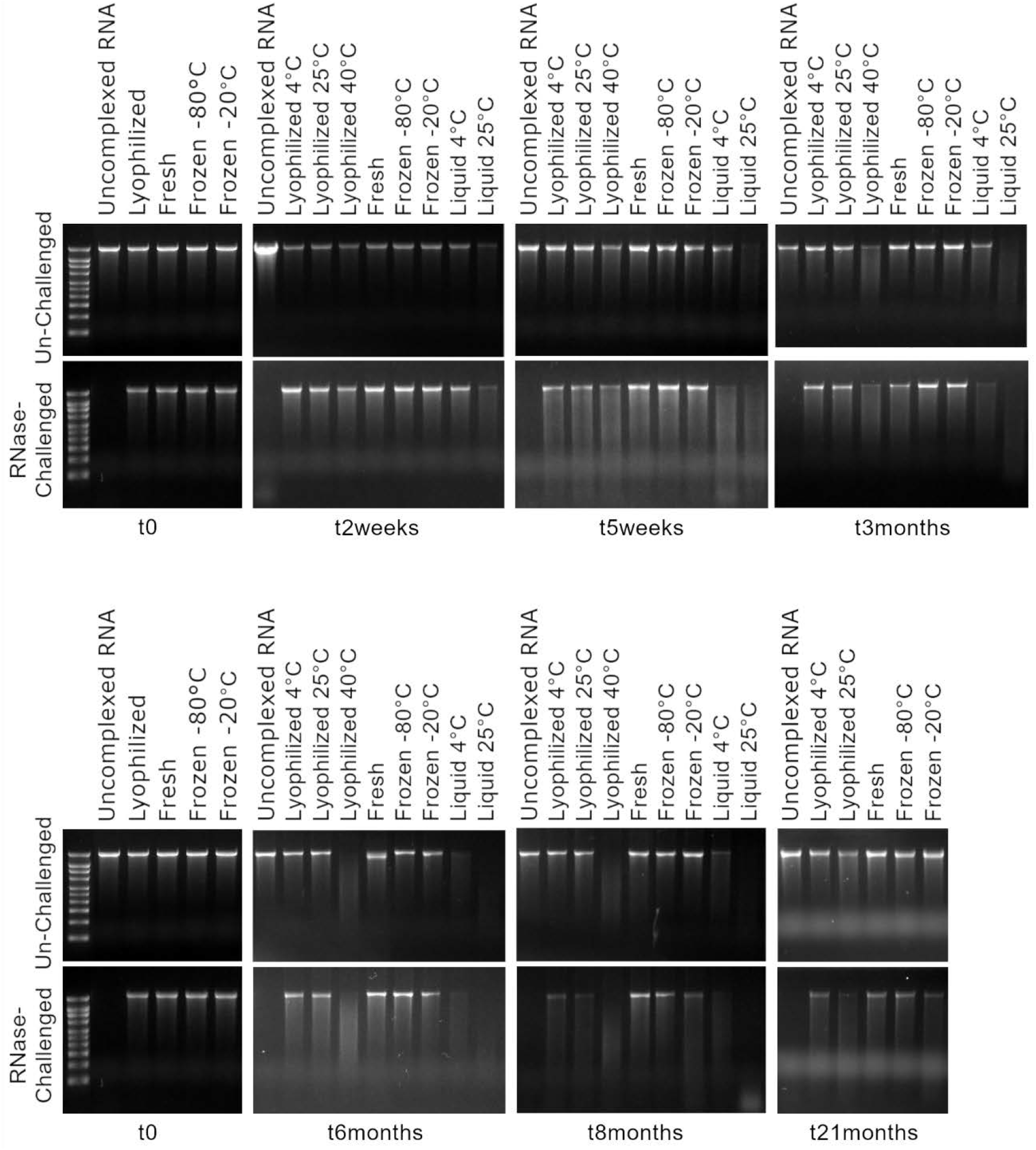
RNA integrity of the stored samples and protection after RNase challenge by agarose gel electrophoresis for each condition at each timepoint.

## References and Notes

1. R. P. Deering, S. Kommareddy, J. B. Ulmer, L. A. Brito, A. J. Geall, Nucleic acid vaccines: prospects for non-viral delivery of mRNA vaccines. Expert Opin Drug Deliv 11, 885–899 (2014).

2. S. Rauch, E. Jasny, K. E. Schmidt, B. Petsch, New Vaccine Technologies to Combat Outbreak Situations. Front Immunol 9, 1963 (2018).

3. C. Zhang, G. Maruggi, H. Shan, J. Li, Advances in mRNA Vaccines for Infectious Diseases. Front Immunol 10, 594 (2019).

4. N. Pardi, M. J. Hogan, F. W. Porter, D. Weissman, mRNA vaccines — a new era in vaccinology. Nature Reviews Drug Discovery 17, 261–279 (2018).

5. L. A. Jackson et al., An mRNA Vaccine against SARS-CoV-2 - Preliminary Report. N Engl J Med 383, 1920–1931 (2020).

6. F. P. Polack et al., Safety and Efficacy of the BNT162b2 mRNA Covid-19 Vaccine. N Engl J Med 383, 2603–2615 (2020).

7. O. S. Kumru et al., Vaccine instability in the cold chain: mechanisms, analysis and formulation strategies. Biologicals 42, 237–259 (2014).

8. D. Chen, D. Zehrung, Desirable attributes of vaccines for deployment in low-resource settings. J Pharm Sci 102, 29–33 (2013).

9. D. J. A. Crommelin, T. J. Anchordoquy, D. B. Volkin, W. Jiskoot, E. Mastrobattista, Addressing the Cold Reality of mRNA Vaccine Stability. Journal of Pharmaceutical Sciences, (2020).

10. U. Sahin, K. Kariko, O. Tureci, mRNA-based therapeutics--developing a new class of drugs. Nat Rev Drug Discov 13, 759–780 (2014).

11. K. A. Whitehead, R. Langer, D. G. Anderson, Knocking down barriers: advances in siRNA delivery. Nat Rev Drug Discov 8, 129–138 (2009).

12. P. S. Kowalski, A. Rudra, L. Miao, D. G. Anderson, Delivering the Messenger: Advances in Technologies for Therapeutic mRNA Delivery. Mol Ther 27, 710–728 (2019).

13. S. Guan, J. Rosenecker, Nanotechnologies in delivery of mRNA therapeutics using nonviral vector-based delivery systems. Gene Ther 24, 133–143 (2017).

14. Y. Y. Tam, S. Chen, P. R. Cullis, Advances in Lipid Nanoparticles for siRNA Delivery. Pharmaceutics 5, 498–507 (2013).

15. Y. Zhao, L. Huang, Lipid nanoparticles for gene delivery. Adv Genet 88, 13–36 (2014).

16. A. M. Reichmuth, M. A. Oberli, A. Jaklenec, R. Langer, D. Blankschtein, mRNA vaccine delivery using lipid nanoparticles. Therapeutic Delivery 7, 319–334 (2016).

17. K. Bahl et al., Preclinical and Clinical Demonstration of Immunogenicity by mRNA Vaccines against H10N8 and H7N9 Influenza Viruses. Mol Ther 25, 1316–1327 (2017).

18. A. M. Reichmuth, M. A. Oberli, A. Jaklenec, R. Langer, D. Blankschtein, mRNA vaccine delivery using lipid nanoparticles. Ther Deliv 7, 319–334 (2016).

19. K. J. Hassett et al., Optimization of Lipid Nanoparticles for Intramuscular Administration of mRNA Vaccines. Mol Ther Nucleic Acids 15, 1–11 (2019).

20. R. L. Ball, P. Bajaj, K. A. Whitehead, Achieving long-term stability of lipid nanoparticles: examining the effect of pH, temperature, and lyophilization. Int J Nanomedicine 12, 305–315 (2017).

21. P. Zhao et al., Long-term storage of lipid-like nanoparticles for mRNA delivery. Bioact Mater 5, 358–363 (2020).

22. L. A. Brito et al., A cationic nanoemulsion for the delivery of next-generation RNA vaccines. Mol Ther 22, 2118–2129 (2014).

23. J. H. Erasmus et al., A Nanostructured Lipid Carrier for Delivery of a Replicating Viral RNA Provides Single, Low-Dose Protection against Zika. Mol Ther 26, 2507–2522 (2018).

24. A. K. Blakney, P. F. McKay, B. I. Yus, Y. Aldon, R. J. Shattock, Inside out: optimization of lipid nanoparticle formulations for exterior complexation and in vivo delivery of saRNA. Gene Ther 26, 363–372 (2019).

25. Materials and methods are available as supplementary materials at the Science website.

26. P. Fonte, S. Reis, B. Sarmento, Facts and evidences on the lyophilization of polymeric nanoparticles for drug delivery. J Control Release 225, 75–86 (2016).

27. K. L. Jones, D. Drane, E. J. Gowans, Long-term storage of DNA-free RNA for use in vaccine studies. Biotechniques 43, 675–681 (2007).

28. B. Petsch et al., Protective efficacy of in vitro synthesized, specific mRNA vaccines against influenza A virus infection. Nat Biotechnol 30, 1210–1216 (2012).

29. M. Alberer et al., Safety and immunogenicity of a mRNA rabies vaccine in healthy adults: an open-label, non-randomised, prospective, first-in-human phase 1 clinical trial. Lancet 390, 1511–1520 (2017).

30. S. Franze, F. Selmin, E. Samaritani, P. Minghetti, F. Cilurzo, Lyophilization of Liposomal Formulations: Still Necessary, Still Challenging. Pharmaceutics 10, (2018).

31. C. Chen, D. Han, C. Cai, X. Tang, An overview of liposome lyophilization and its future potential. J Control Release 142, 299–311 (2010).

32. C. Liu et al., Barriers and Strategies of Cationic Liposomes for Cancer Gene Therapy. Mol Ther Methods Clin Dev 18, 751–764 (2020).

